# In vivo itaconate tracing reveals degradation pathway and turnover kinetics

**DOI:** 10.1101/2025.01.02.629617

**Authors:** Hanna F. Willenbockel, Alexander T. Williams, Alfredo Lucas, Birte Dowerg, Pedro Cabrales, Christian M. Metallo, Thekla Cordes

## Abstract

Itaconate is an immunomodulatory metabolite that alters mitochondrial metabolism and immune cell function. This organic acid is endogenously synthesized via tricarboxylic acid (TCA) metabolism downstream of TLR signaling. Itaconate-based treatment strategies are being explored to mitigate numerous inflammatory conditions. However, little is known about the turnover rate of itaconate in circulation, the kinetics of its degradation, and the broader consequences on metabolism. By combining mass spectrometry and in vivo ^13^C itaconate tracing, we demonstrate that itaconate is rapidly eliminated from plasma, excreted via urine, and fuels TCA cycle metabolism specifically in the liver and kidneys. These studies further revealed that itaconate is converted into acetyl-CoA, mesaconate, and citramalate in mitochondria. Itaconate administration also influenced branched-chain amino acid metabolism and succinate levels, indicating a functional impact on succinate dehydrogenase (SDH) and methylmalonyl-CoA mutase (MUT) activity. Our findings uncovered a previously unknown aspect of the itaconate metabolism, highlighting its rapid catabolism in vivo that contrasts findings in cultured cells.

## Introduction

The small molecule itaconate is endogenously synthesized by immune cells, and the dynamic control of itaconate modulates metabolic pathways and immune responses^1–3^. Itaconate functions as a competitive inhibitor of succinate dehydrogenase (SDH), thereby influencing mitochondrial metabolism and immune cell function^4,5^. Despite extensive research on the biosynthetic pathway catalyzed by aconitate decarboxylase (ACOD1), also known as immune-responsive gene 1 protein (IRG1), the itaconate degradation pathway remains poorly understood^6,7^. Itaconate is metabolized to itaconyl-coenzyme A (CoA)^8–10^, mesaconate^11–13^, and citramalyl-CoA^9,11^. Further, degradation into pyruvate and acetyl-CoA has been documented in liver mitochondria in the early 1960s^14,15^. However, in prior ^13^C itaconate tracer studies we were unable to detect labeling on TCA cycle intermediates in cultured mammalian cell models including those of the brain, immune system, and hepatocytes indicating that itaconate is not metabolized to acetyl-CoA in cultured cells^11,16,17^. Itaconate has been applied therapeutically to mitigate the consequences of inflammatory stress in vivo^16,18,19^. However, the consequences of high circulating concentrations and the kinetics of its degradation in vivo remain unclear. To address this limitation, we performed ^13^C itaconate tracing in vivo and quantified itaconate dissimilation pathways and turnover rates.

## Main

### Dynamic itaconate metabolism and correlation with succinate levels in infused rat models

Itaconate accumulates at high levels in cells, but levels detected in circulation are in the µM range^20^. To increase systemic itaconate levels, we established in our previous study an itaconate treatment strategy in an animal model of ischemia-reperfusion (I/R) and demonstrated that itaconate mitigates cellular injuries associated with reoxygenation^16^. We observed that itaconate was rapidly cleared from plasma and introduced a model in which itaconate transiently inhibits SDH activity to mitigate reperfusion-induced overactivation of SDH. Since itaconate-induced SDH inhibition is reversible itaconate levels might be fine-tuned to modulate cellular metabolism and function^16^. To gain more quantitative insights into the dynamics of itaconate metabolism, we infused rat animal models with 15 mg/kg/min itaconate for 30 min and subsequently quantified plasma metabolite levels over time (Extended Data Fig. 1a). Infusion with itaconate for 30 min significantly elevated plasma concentrations of the TCA cycle intermediates succinate and malate, while other metabolites were less affected (Fig. 1a). This data indicate that the TCA cycle metabolism might be the primary target of short term itaconate treatments by modulating SDH activity. Next, we conducted a time-series analysis of plasma metabolite levels during two itaconate infusions with each 30 min (Extended Data Fig. 1a). The plasma concentration of itaconate increased to approximately 0.45 mM and most itaconate was cleared within 60 min. The second administration of itaconate yielded comparable outcomes indicating that in vivo itaconate metabolism is highly dynamic (Fig. 1b). Next, we calculated the pharmacokinetic parameters for itaconate and observed an elimination half-time (T_½_) of 53 min following the initial infusion and 85 min following the second infusion (Fig. 1b, Extended Data Fig.1b). Our kinetic parameters indicate that itaconate is rapidly cleared in vivo suggesting a dynamic and reversible impact of itaconate on metabolism. Indeed, we observed that succinate concentrations correlated strongly with itaconate levels in the plasma indicating reversible SDH activity inhibition (Fig. 1c). Itaconate also affected abundances of other TCA cycle intermediates, specifically malate suggesting that itaconate-induced SDH inhibition may have additional effects on TCA cycle metabolism and related amino acids (Extended Data Fig. 1c-e).

**Figure 1:**
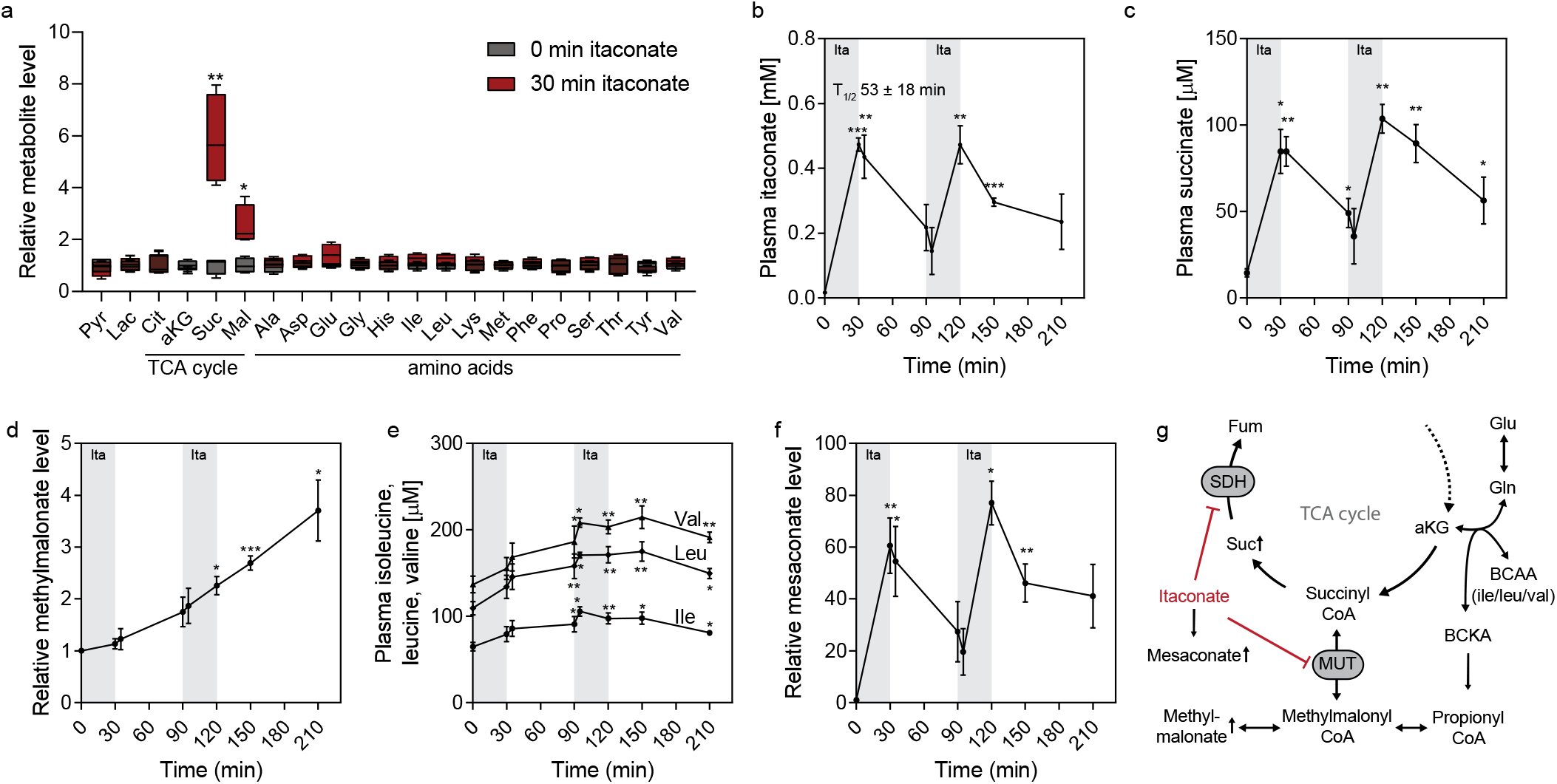
Dynamic itaconate metabolism and correlation with succinate levels in infused rat models. **a**, Plasma metabolite abundances at 30 min compared to 0 min after itaconate infusion. **b**, Plasma itaconate levels over time with two itaconate infusions each for 30 min as indicated in grey. **c**, Plasma succinate levels over time with two itaconate infusions. **d**, Relative level of methylmalonate in plasma over time with two itaconate infusions. **e** Plasma level of leucine, isoleucine, and valine over time with two itaconate infusions. **f**, Relative mesaconate level over time with two itaconate infusions. **g**, Schematic depicting the impact of itaconate on succinate dehydrogenase (SDH) and mutase (MUT) activities. Experiments were performed with n = 4 rats and itaconate was infused twice for each 30 min with 15 mg/kg/min indicated in grey. Data are presented as box (25^th^ to 75^th^ percentile with median line) and whiskers (min. to max. values) (a) or means ± s.e.m. (b-f) obtained from n = 4 rats. *P* values were calculated by multiple unpaired *t*-test (a) or two-way *ANOVA* compared to 0 min (b-f) with * *p* < 0.05; ** *p* < 0.01; *** *p* < 0.001.

Itaconate is metabolized to itaconyl-CoA, an inhibitor of both methylmalonyl-CoA mutase (MUT) and aminolevulinate synthase (ALAS) influencing mitochondrial coenzyme B_12_ and heme metabolism^8–10^. Impaired MUT activity leads to elevated levels of methylmalonate (MMA) and altered branched-chain amino acid (BCAA) metabolism^8,9,17^. In contrast to succinate levels, plasma MMA levels demonstrated a gradual accumulation over time indicating itaconate-induced MUT inactivation (Fig. 1d). Furthermore, we observed that the levels of BCAAs leucine, isoleucine, and valine were significantly increased over time in response to itaconate treatments (Fig. 1e, Extended Data Fig. 1f-h). This data suggests that the impact on BCAA metabolism may be less dynamic compared to succinate levels. Our observation further supports our previous in vitro studies indicating that itaconate-derived itaconyl-CoA irreversibly inactivates mitochondrial MUT activity leading to increased MMA levels and altered BCAA metabolism^8,9,17^. In contrast to BCAAs, other plasma amino acids were less affected over time in response to itaconate (Extended Data Fig. 1i-t). Moreover, we observed a significant correlation between plasma mesaconate, a metabolite synthesized from itaconate, levels and those of itaconate and succinate (Fig. 1b, c, f)^11,13^. Our study revealed that itaconate is rapidly cleared in vivo and consequently impacts mitochondrial and BCAA metabolism indicating altered SDH and MUT activities (Fig. 1g).

### Itaconate is a substrate for mitochondrial metabolism in mouse liver and kidney

Since itaconate is rapidly cleared from plasma (Fig. 1b), we postulated that itaconate might be taken up and further metabolized by tissues. To better understand the fate of itaconate, we applied a ^13^C itaconate tracer and quantified itaconate fluxes in vivo. Specifically, we administered a dose of 400 mg/kg body weight [U-^13^C_5_]itaconate to mice and quantified metabolite abundances and labeling over time on plasma and tissue metabolome (Extended Data Fig. 2a). We observed that 15 min after itaconate administration, plasma itaconate levels increased to about 2.5 mM and were cleared within 45 min with T_½_ = 10.9 min (Fig. 1a, Extended Data Fig. 2b). We also observed a robust positive correlation between succinate and itaconate plasma levels in mice over time (Fig. 2b) indicating functional involvement of SDH activity. The abundances of other TCA cycle intermediates, such as malate, also correlated with itaconate levels indicating a dynamic impact of itaconate on mitochondrial TCA cycle metabolism as previously observed in our rat model (Fig. 2c, Fig. 1). Further, we quantified a potential impact on MUT activity and observed that levels of MMA and BCAAs in the plasma were not affected after short time upon itaconate treatments (Extended Data Fig. 3a, b). However, MMA and BCAAs increased in some tissues indicating that itaconate may affect Vitamin B_12_ metabolism and MUT activity in the liver and the heart, as previously observed in our rat model with longer itaconate treatments (Extended Data Fig. 3c-f, Fig. 1).

**Figure 2:**
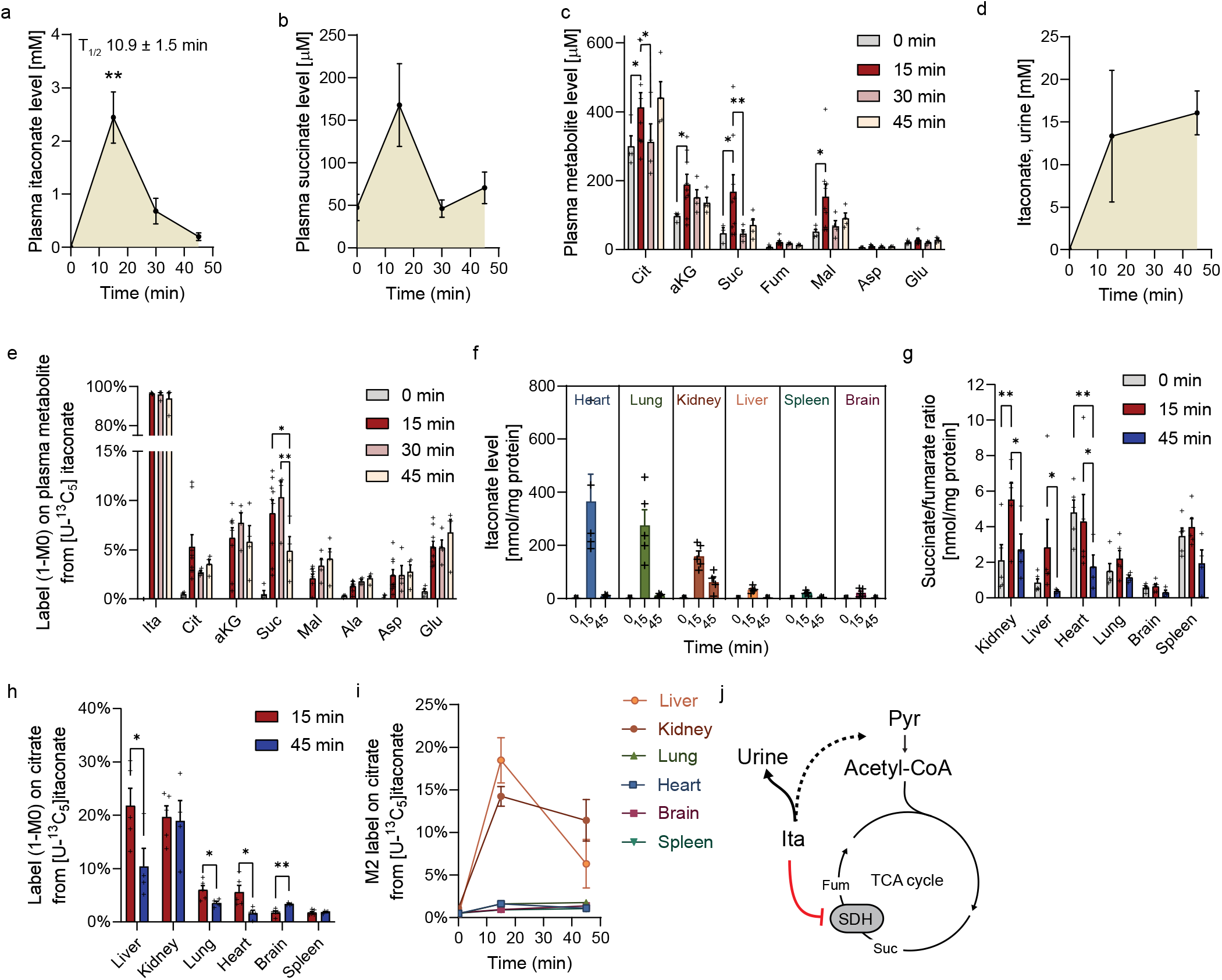
Itaconate is a substrate for mitochondrial metabolism in mouse liver and kidney. **a**, Plasma itaconate levels in response to itaconate treatment with elimination halftime (T_1/2_). **b**, Plasma succinate levels in response to itaconate treatment. **c**, Levels of TCA cycle intermediates after itaconate treatment for indicated times. **d**, Itaconate levels in urine. **e**, Labeling (1 - M0) on TCA cycle intermediates in plasma from [U-^13^C_5_]itaconate. **f**, Levels of itaconate in different tissues. **g**, Ratio of succinate over fumarate after itaconate treatment in different tissues. **h**, Labeling (1 - M0) on citrate from [U-^13^C_5_]itaconate in different tissues. **i**, M2 labeling on citrate from [U-^13^C_5_]itaconate in different tissues. **j**, Schematic depicting itaconate clearance pathway and succinate dehydrogenase (SDH) inhibition. Stashed arrows depict indirect metabolic pathways. Mice were injected with 400 mg/kg body weight [U-^13^C_5_]itaconate. Data are presented as means ± s.e.m. Plasma samples at 0 min (n = 4), 15 min (n = 9), 30 min (n = 4), and 45 min (n = 4); tissue samples at 0 min (n = 4), 15 min (n = 5), and 45 min (n = 4); urine samples at 0 min (n = 3), 15 min (n = 4), and 45 min (n = 3). One-way *ANOVA* compared to 0 min itaconate (a, b, d), or two-way *ANOVA* (c, e-h) with * *p* < 0.05; ** *p* < 0.01, *** *p* < 0.001, ^#^ *p* < 0.0001.

Previous studies demonstrated that when itaconate was fed to dogs, approximately 25 % of the itaconate could be recovered in the urine suggesting that parts of itaconate might be metabolized by tissues^21^. Therefore, we elucidated potential clearance pathways for ^13^C itaconate in vivo and observed approximately 15 mM itaconate in the urine (Fig. 2d). These results suggest that renal excretion is a major pathway for itaconate clearance. We also observed dynamic labeling on plasma TCA cycle intermediates and related amino acids 15 min after tracer injection indicating that itaconate is metabolized further and used for mitochondrial TCA cycle metabolism (Fig. 2e). To identify the impact of circulating itaconate on tissue metabolism and identify potential tissues involved in itaconate dissimilation, we quantified levels and labeling on TCA cycle-related metabolites in diverse tissues. Itaconate was detected in all tissues after 15 min, with the highest concentrations observed in the kidney, heart, and lung, and lower levels in the liver, brain, and spleen (Fig. 2f). Though a moderate increase in succinate was observed in tissues, succinate to fumarate ratios in the kidney and liver were significantly elevated indicating impaired SDH activity (Fig. 2g, Extended Data Fig. 2c-j). Itaconate labeled around 20 % of the liver and kidney citrate pool and up to 10 % in other tissues (Fig. 2h). Notably, the liver and kidney appear to be highly involved in the itaconate dissimilation pathway, as M2 labeling on citrate reached approximately 20 %, while citrate labeling in other tissues remained below 5 % (Fig. 2i). Collectively, this data indicates that itaconate is cleared from plasma by renal clearance and metabolized via mitochondrial metabolism in the liver and kidney (Fig. 2j).

### The fate of ^13^C itaconate in liver tissue

Next, we quantified mass isotopomer distributions on TCA cycle intermediates from [U-^13^C_5_]itaconate to identify potential pathways involved in itaconate dissimilation. Given that citrate was the most heavily labeled metabolite in the liver tissue, we monitored the ^13^C-derived itaconate carbons in this tissue at 15 and 45 min after itaconate injection (Fig. 2i). We observed that the amount of fully labeled M5 itaconate was nearly 100 % indicating that the endogenous itaconate present in the tissue was negligible (Fig. 3a). While the levels of pyruvate, lactate, and alanine exhibited lower labeling, approximately 20 % of M2 citrate was labeled (Fig. 3b-e). Further, additional TCA cycle intermediates were robustly labeled with M2 including -ketoglutarate, fumarate, succinate, and malate, as well as related amino acids, glutamate, and aspartate (Fig. 3f-k). These labeling patterns indicate that itaconate is not converted to citrate via the reverse ACOD1 synthesis pathway which would result in M5 on citrate. Conversely, itaconate is metabolized to a labeled C_2_ compound, which fuels the citrate pool, resulting in an M2 labeling on citrate (Fig. 3e). We quantified high itaconyl-CoA abundances in itaconate-treated liver tissue while no itaconyl-CoA was detectable in control conditions indicating that itaconyl-CoA is an intermediate of the itaconate dissimilation pathway (Fig. 3l). Given the small amount of labeling observed on M3 pyruvate, we proceeded to quantify acetyl-CoA levels, which demonstrated a modest increase in response to itaconate treatment (Fig. 3l). Further, we detected elevated levels of acetyl-carnitine suggesting that the carnitine shuttle may be involved in the itaconate dissimilation pathway (Fig. 3l). ^13^C itaconate highly labeled itaconyl-CoA, approximately 20 % of the acetyl-CoA pool, and comparable labeling was observed on acetyl-carnitine suggesting that itaconate-derived carbons are transported into the mitochondria via the carnitine shuttle (Fig. 3m). These carbons may be subsequently utilized to form M2 labeling on citrate and other TCA cycle intermediates via the acetyl-CoA pathway. Thus, itaconate may serve as a carbon fuel for TCA cycle metabolism via acetyl-CoA resulting in M2 labeling on citrate specifically in the kidney and liver (Fig. 2i). Indeed, the mass isotopologue distribution of TCA cycle intermediates in the kidney revealed the presence of M2 labeling on TCA cycle intermediates, while pyruvate, lactate, and alanine exhibited M3 labeling derived from ^13^C itaconate (Extended Data Fig. 4). Thus, itaconate may fuel TCA cycle metabolism via acetyl-CoA, particularly in the liver and kidney.

**Figure 3:**
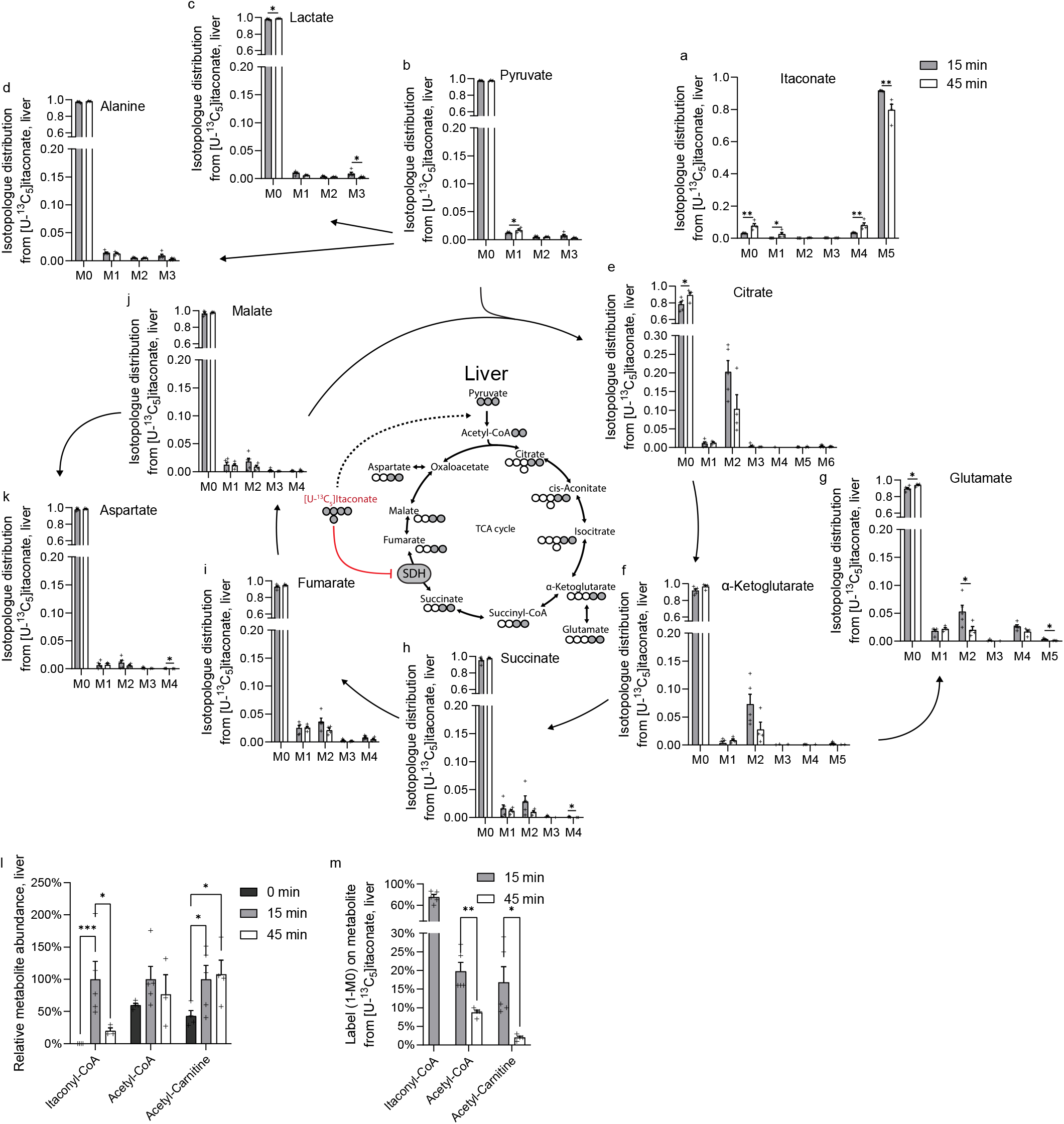
The fate of ^13^C itaconate in liver tissue. **a-k**, Isotopologue distribution from [U-^13^C_5_]itaconate on **a**, itaconate, **b**, pyruvate, **c**, lactate, **d**, alanine, **e**, citrate, **f**, -ketoglutarate, **g**, glutamate, **h**, succinate, **i**, fumarate, **j**, malate, and **k**, aspartate in liver tissue. **l**, Abundances of itaconyl-CoA, acetyl-CoA, and acetyl-carnitine relative to 15 min baseline corrected to 100 %. **m**, Labeling (1 - M0) on itaconyl-CoA, acetyl-CoA, and acetyl-carnitine from [U-^13^C_5_]itaconate. Schematic of [U-^13^C_5_]itaconate used for TCA cycle metabolism with open bars depicting ^12^C and closed circles ^13^C shown in the middle of the figure. Mice were injected with 400 mg/kg body weight [U-^13^C_5_]itaconate. Data are presented as means ± s.e.m with tissue samples at 0 min (n = 4), 15 min (n = 5), and 45 min (n = 4). Multiple unpaired *t*-test (a-k, m) or two-way *ANOVA* (l) with * *p* < 0.05; ** *p* < 0.01, *** *p* < 0.001.

### Itaconate is metabolized to mesaconate and citramalate in vivo

In our previous studies, we traced various cell types, including immune cells, neurons, astrocytes, and hepatocarcinoma cells, with ^13^C itaconate but did not detect labeling on citrate^11,16,17^. Since these studies were performed with 2 mM or lower concentrations of ^13^C itaconate, we cultured the hepatocarcinoma cell lines HepG2 and Huh7 in the presence of 1 mM and 10 mM [U-^13^C_5_]itaconate for 24 h. Similar to our previous studies, we did not observe labeled citrate or palmitate, even after treatment with 10 mM ^13^C itaconate. This data suggests that itaconate is not metabolized to acetyl-CoA in vitro, which serves as a fuel for the TCA cycle metabolism and de novo lipogenesis (Fig. 4a). Thus, the itaconate dissimilation pathway may exhibit differential behavior in cells compared to in vivo conditions. Subsequently, we quantified the labeling of mesaconate and citramalate, two metabolites derived from itaconate^11,14,15,22^. We observed fully M5 labeled mesaconate and citramalate in both cell lines indicating that all five carbons for the carbon backbone were derived from itaconate (Fig. 4a, Extended Data Fig. 5a, b). Since our mice studies were conducted with more short time points we also quantified time-dependent metabolic fluxes and traced cells for 15, 30, and 60 min. Labeling of citrate was negligible, while mesaconate, citramalate, and itaconate were fully labeled even at the 15 min time point (Fig. 4b, Extended Data Fig. 5c). We also observed that parts of the labeled mesaconate, citramalate, and citrate were present in the urine indicating renal clearance (Extended Data Fig. 5d). Thus, itaconate provides the five-carbon backbone for the formation of mesaconate and citramalate, while itaconate-derived carbons also fueled TCA cycle metabolism in our in vivo models (Extended Data Fig. 5e).

**Figure 4:**
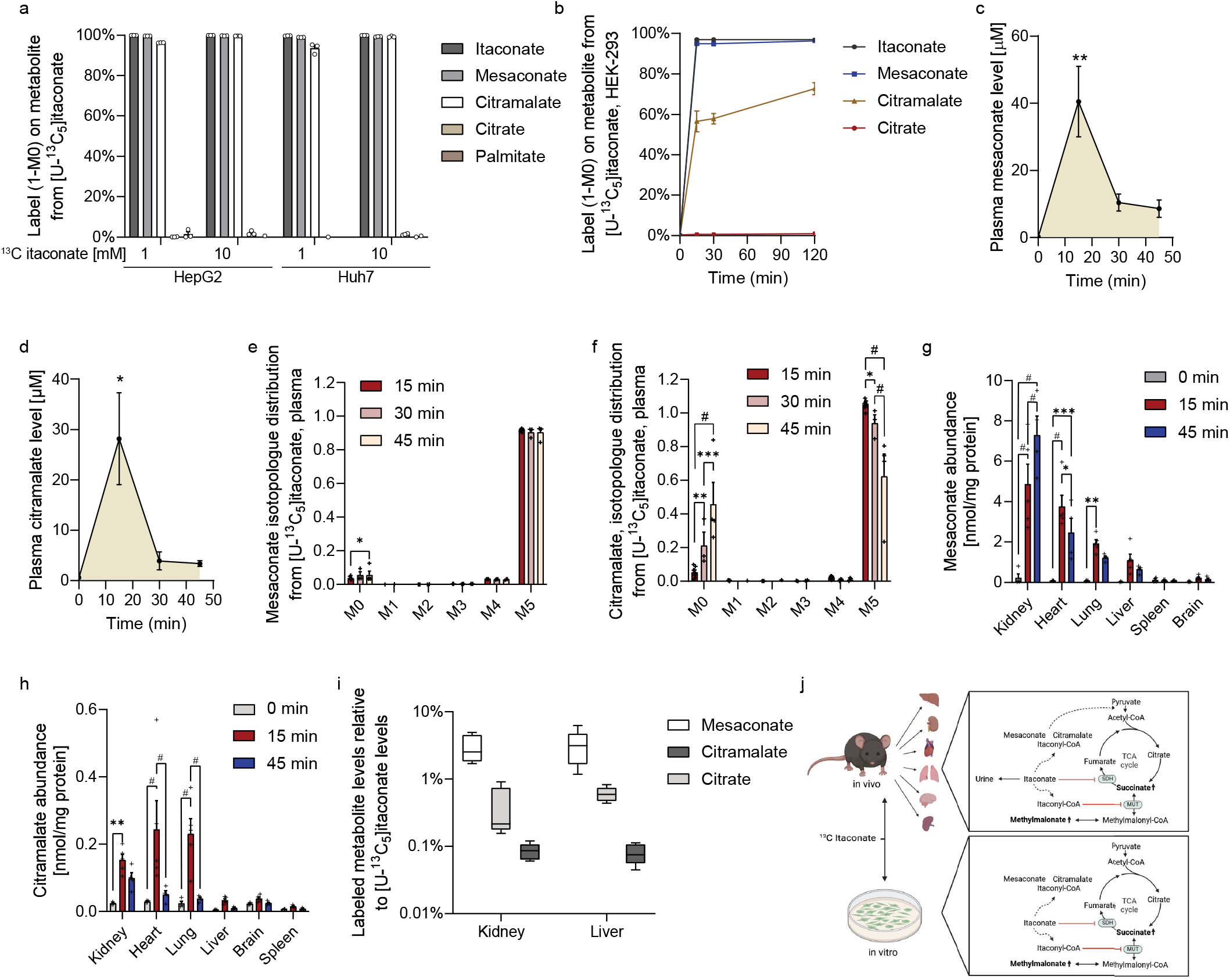
Itaconate is metabolized to mesaconate and citramalate in vivo. **a**, Labeling (1 – M0) on itaconate, mesaconate, citramalate, citrate, and palmitate in HepG2 and Huh7 cells after 24 h culture with 1 mM and 10 mM [U-^13^C_5_]itaconate. **b**, Labeling (1 – M0) on itaconate, mesaconate, citramalate, and citrate in HEK-293 cells cultured for 15, 30, and 120 min with 3 mM [U-^13^C_5_]itaconate. **c**, Plasma mesaconate level. **d**, Plasma citramalate level. **e**, Plasma mesaconate isotopologue distribution, **f**, Plasma citramalate isotopologue distribution. **g**, Labeled mesaconate abundance in tissues. **h**, Labeled citramalate abundance in tissues. **i**, Labeled metabolite relative to itaconate levels in liver and kidney tissues 15 min after ^13^C itaconate injection. **j**, Schematic depicting comparing potential itaconate dissimilation in *in vivo* and *in vitro* models. Mice were injected with 400 mg/kg body weight [U-^13^C_5_]itaconate. Data are presented as means ± s.e.m (a-h) or box (25^th^ to 75^th^ percentile with median line) and whiskers (min. to max. values) (i). Plasma samples at 0 min (n = 4), 15 min (n = 9), 30 min (n = 4), and 45 min (n = 4); tissue samples at 0 min (n = 4), 15 min (n = 5). One-way *ANOVA* compared to 0 min itaconate (c, d), or two-way *ANOVA* (e-i) with * *p* < 0.05; ** *p* < 0.01, *** *p* < 0.001, ^#^ *p* < 0.0001.

Given that mesaconate and citramalate are derived from itaconate, we quantified the kinetics of these C_5_ dicarboxylate compounds in vivo following itaconate treatment. Our study revealed a robust correlation between plasma levels of mesaconate and citramalate and those of itaconate (Fig. 4c, d). Mesaconate labeling indicates that it is directly derived from itaconate and negligible endogenous mesaconate was present in these otherwise healthy animals (Fig. 4e, Extended Data Fig. 5f, g). In contrast, citramalate was also M5 labeled, yet the labeling decreased over time indicating dynamic itaconate metabolism and the presence of endogenous, unlabeled citramalate (Fig. 4f, Extended Data Fig. 5h). Further, mesaconate and citramalate were synthesized in a variety of tissues, with kidney showing the highest abundances (Fig. 4g). To gain further insight into the extent of itaconate metabolism to mesaconate, citramalate, and citrate, we normalized the labeled abundances of these metabolites to labeled itaconate abundance at 15 minutes post-injection present in kidney and liver tissues. We observed that approximately 5 % of itaconate is converted to mesaconate, 1 % to citrate, and below 1 % to citramalate (Fig. 4i). Thus, circulating itaconate is excreted via renal clearance and a small fraction undergoes further metabolism with involvement of kidney and liver. Further, C_5_ dicarboxylate compounds mesaconate and citramalate are derived from itaconate in both in vivo and in vitro conditions whereas conversion into citrate occurred in in vivo conditions only (Fig. 4j).

## Discussion

Our ^13^C tracing study elucidated the turnover kinetics and dissimilation pathways to provide new insights into the dynamic itaconate metabolism, pharmacokinetics, and effects on TCA cycle and BCAA metabolism through the inhibition of SDH and MUT activity. We identified renal excretion as the major pathway for itaconate clearance, with a small fraction of itaconate being metabolized to acetyl-CoA, which fuels TCA cycle metabolism, or converted to mesaconate and citramalate. This metabolic fate of itaconate suggests a previously unidentified metabolic function for itaconate, linking itaconate to substrate utilization and metabolic regulation.

Our in vivo study demonstrates that itaconate levels correlate strongly with plasma succinate levels suggesting a regulatory role for itaconate in systemic succinate levels and subsequent immune responses^23^. Since SDH is complex II of the mitochondrial respiratory chain, this dynamic regulation of SDH may be beneficial in situations of limited oxygen availability, such as diseases associated with reoxygenation injury. Our in vivo data support the model that itaconate functions as a competitive and reversible SDH inhibitor, which gradually awakens mitochondrial flux and thereby mitigates reperfusion injury^5,16,17^. While single doses of itaconate rapidly affected SDH function, effects on BCAA metabolism and methylmalonate were less reversible, potentially due to MUT inactivation via itaconyl-CoA^8,17^. These findings further support our previous observation in cultured cell models indicating that itaconate may affect transaminase reactions^17^. The long-term effects on BCAA and vitamin B_12_ metabolism may have implications in cell types with highly activated BCAA metabolism, such as adipocytes^24,25^.

Our tracing study reveals significant effects of itaconate on the TCA cycle and CoA metabolism in the liver. These findings are consistent with previous research demonstrating that itaconate exerts beneficial effects in metabolic disorders including diet-induced obesity ^18,26^ and non-alcoholic fatty liver disease^27^. Itaconate is transported in the liver via the sodium dicarboxylate cotransporter^26,28^. Thus, circulating itaconate may impact hepatic metabolism and our itaconate clearance data may provide insight into potential treatment strategies. Due to its water-soluble nature, the addition of itaconate to drinking water^29^ or fluid therapies^16^ may facilitate long-term and continuous administration of itaconate. Further research is needed to elucidate the long-term effects of itaconate supplementation and optimize dosing regimens.

Itaconate has antimicrobial properties and certain bacterial species have developed strategies to use itaconate as a carbon source for detoxification^30^. Thus, the itaconate dissimilation pathway may be conserved across species and further research is needed to elucidate the discrepancies of itaconate dissimilation pathway observed in different model systems. Additional investigation is also needed to identify potential metabolites and enzymes involved in this pathway. For example, enzymes involved in the synthesis of itaconyl-CoA, such as succinyl-CoA:glutarate-CoA transferase (SUGCT)^10^ and succinyl-CoA synthetase^14,15^ may influence substrate phosphorylation and CoA homeostasis^12,17^. While numerous studies have focused on the immunomodulatory effects of itaconate, our present study provides a novel perspective on its role in regulating metabolism^3,31–33^. Itaconate also possesses beneficial properties in the context of cancer, obesity, and other diseases^18,28,34–37^. Thus, some effects of itaconate may be attributed to itaconate degradation products and subsequent metabolic reprogramming that may be beneficial in diverse disease settings.

## List of abbreviation

^12^C: Carbon-12
^13^C: Carbon-13
[U-^13^C_5_]itaconate: Uniformly ^13^C labeled itaconate
ACOD1: Aconitate decarboxylase 1
ALAS: Aminovlevulinate synthase
AUC: Area under the curve
BCAA: Branched-chain amino acid
C_max_: Maximal plasma concentration
CoA: Coenzyme-A
FAMES: Fatty acid methyl esters
GC: Gas chromatograph
I/R: ischemia-reperfusion
IRG1: Immune responsive gene 1 (protein)
*Irg1*: Immunoresponsive gene 1 (gene)
LC: Liquid chromatography
MMA: Methylmalonate
MS: Mass spectrometry
MUT: Methylmalonyl-CoA mutase
SDH: Succinate dehydrogenase
SUGCT: Succinyl-CoA:glutarate-CoA transferase
T_1/2_: Elimination halftime
TCA: Tricarboxylic acid

## Material and Methods

### Animal studies

Animal handling and care followed the NIH Guide for Care and Use of Laboratory Animals (Protocols: S00149 and S11306). The experimental protocol was approved by the UCSD Institutional Animal Care and Use Committee.

### Itaconate infusion experiments in rat models

Itaconate infusion studies were performed in male Sprague-Dawley rats (Harlan Laboratories, Indianapolis, Ind) weighing 200 to 250 g. Animals were anesthetized with isoflurane in compressed room air (Drägerwerk AG, Lübeck, Germany) and placed on a heating pad to maintain core body temperature at 37 °C for the duration of the experiment. A femora catheter was implanted and itaconate was infused with 15 mg/kg/min for 30 min. The infusion was stopped for 60 min and itaconate was then infused again for 30 min for a second round. Plasma samples were taken at timepoints as indicated in the text with n = 4 animals. Data for rat infusion experiments are depicted in Fig. 1 and Extended Data Fig. 1.

### ^13^C itaconate study in mouse models

C57BL/6J mice were obtained from Jackson Laboratories (Bar Harbor, ME). Mice were administered 400 mg/kg body weight itaconate via retroorbital injection^38^. Itaconate was prepared in NaCl and adjusted to pH = 7.3. Mice were fasted for 6 h before itaconate injection and the age of male mice was 8 weeks. Plasma and tissues were collected at indicated time points (Extended Data Fig. 2a) and further used for metabolite analysis. Animals were anesthetized with isoflurane, decapitated, and tissues rapidly collected, frozen to temperatures of liquid nitrogen, and stored at −80 °C until analysis. Tissues were collected from n = 5 animals for 15 min ^13^C itaconate and n = 4 for 45 min ^13^C itaconate treatment. Data was compared to the control condition (0 min itaconate) where NaCl was given for 45 min with n = 4 animals.

### Cell culture

The cell lines HEK-293 (ATTC, CRL-1573), HepG2 (ATCC HB-8065), and Huh7 (provided by M. Hermann, MIT, Cambridge, MA, USA) were used for the experiments. Cells were tested negative for mycoplasma contamination by MycoAlert® Mycoplasma Detection Kit (Lonza). Cells were cultured in Dulbecco’s Modified Eagle Media (DMEM, Cat. #11965-092, Gibco) containing 25 mM glucose, 4 mM glutamine, 100 U/ml penicillin, and 100 µg/ml streptomycin in a humidified cell culture incubator at 37 °C and 5 % CO_2_. Medium was supplemented with 10 % fetal bovine serum (FBS) (Cat. #16000-044, Gibco) and cells were detached with 0.05 % trypsin-EDTA.

### Gas chromatograph - Mass spectrometry (GC-MS), sample preparation, and analysis

Metabolites were extracted, analyzed, and quantified, as previously described in detail^39^. Briefly, plasma metabolite levels were extracted using 10 µl plasma and 90 µl methanol:water (8:1). Urine metabolites were extracted using 5 µl plasma and 45 µl methanol:water (8:1). Tissues were pulverized using a cellcrusher cryogenic tissue pulverizer (Cellcrusher, Cork, Ireland). The powder was stored at -80 °C until further usage. 10 - 20 mg pulverized tissue was homogenized with a ball mill (Retsch Mixer Mill MM 400) at 30 Hz for 3 min and metabolites were extracted with 0.5 ml -20 °C methanol, 0.2 ml 4°C cold water, and 0.5 ml -20 °C chloroform. 50 µl was taken before chloroform addition to determine tissue protein content using BCA protein assay (Lambda Biotech, G1002) for normalization. For absolute quantification of amino acids (Fig. 1, Extended Data Fig. 1), 10 µl of 100 µM labeled (^13^C,^15^N) amino acid standard mix (MSK-A2-1.2., Cambridge Isotope Laboratories, Inc) was spiked to each sample (1 nmoles per sample). Other metabolites including TCA cycle intermediates were quantified based on external standard curves. The plasma and tissue extracts were vortexed for 10 min at 4 °C and centrifuged at 16,000 × *g* for 5 min at 4 °C. The upper aqueous phase was evaporated under vacuum at 4 °C. Derivatization for polar metabolites was performed using a Gerstel MPS with 15 µl of 2 % (w/v) methoxyamine hydrochloride (Thermo Scientific) in pyridine (incubated for 60 min at 45 °C) and 15 µl N-tertbutyldimethylsilyl-N-methyltrifluoroacetamide (MTBSTFA) with 1 % tert-butyldimethylchlorosilane (Regis Technologies) (incubated further for 30 min at 45 °C). Derivatives were analyzed by GC-MS using a DB-35MSUI column (30 m × 0.25 i.d. × 0.25 µm) installed in an Agilent 7890B gas chromatograph (GC) interfaced with an Agilent 5977A mass spectrometer (MS) operating under electron impact ionization at 70 eV. The MS source was held at 230 °C and the quadrupole at 150 °C and helium was used as carrier gas. The GC oven was held at 100 °C for 2 min, increased to 300 °C at 10 °C/min, and held at 325 °C for 3 min.

The lower organic phase from Huh7 and HepG2 culture cells were derivatized to form fatty acid methyl esters (FAMES) using 500 pl 2% H_2_SO_4_ in MeOH and incubation at 50 °C for 2 h. FAMES were extracted via the addition of 100 µl saturated salt solution and 500 µl hexane. FAMES were analyzed using a Select FAME column (100 m × 0.25 mm i.d.) installed in an Agilent 7890 A GC interfaced with an Agilent 5975C MS. Helium was used as a carrier gas and the GC oven was held at 80 °C, increased by 20 °C/min to 170 °C, increased by 1 °C/min to 204 °C, then 20 °C/min to 250 °C and hold for 10 min.

### Measurements of CoA and carnitine species

Itaconyl-CoA, acetyl-CoA, and acetyl-carnitine were measured using reversed-phase liquid chromatography (RPLC) method, as described in our previous publication^17^. Briefly, 10 mg pulverized tissue was homogenized with a ball mill (Retsch Mixer Mill MM 400) at 30 Hz for 5 min. Metabolites were extracted with 1 ml −20 °C 80 % methanol/water and the extracts were centrifuged at 16,000 × *g* for 5 min at 4 °C. 200 µl of the extracts were dried under airflow, resuspended in 100 l Buffer A and 5 µl of the sample was measured on a liquid chromatography (LC) coupled to a Q Exactive system (Q Exactive Hybrid Quadrupole-Orbitrab MS w/Vanquish Flex Binary UHPLC system, Thermo Scientific, Waltham, MA, USA). A C18 column (C18 1.7 µm, 100Å, 100 × 2.1 mm, Phenomenex, Cat.#00D-4475-AN) was employed with mobile phase Buffer A (5 mM ammonium acetate in water, pH = 6.8) and Buffer B (100 % methanol). The QE-MS was operated in positive mode. Metabolites were verified with external standards or specific MS2 fragments. Mass accuracy obtained for all metabolites was below 5 ppm. Data were acquired with Thermo Xcalibur software and analyzed using EL-Maven software with correction for natural abundance^40^. Abundance was normalized to mg tissue. Itaconyl-CoA was detected in itaconate-treated samples only and was M5 labeled from [U-^13^C_5_]itaconate.

### Isotopic tracing and analysis

Labeled [U-^13^C_5_]itaconate was provided by the Metabolite Standards Synthesis Core, arranged through the National Institutes of Health Common Fund’s Metabolomics Initiative^41^. Cells were cultured in growth medium containing 10 % FBS and ^13^C itaconate. All media was adjusted to pH = 7.3. Huh7 and HepG2 cells were cultured for 24 h in the presence of 1 mM and 10 mM ^13^C itaconate. HEK-293 cells were cultured for 15, 30, and 120 min with 3 mM ^13^C itaconate. Metabolites were extracted and labeling on metabolites from ^13^C itaconate was quantified using GC-MS technology. Mass isotopomer distributions and total metabolite abundances were computed by integrating mass fragments using a MATLAB based algorithm with corrections for natural isotope abundances as described previously^39,42^. Labeling is depicted as 1-M0 or isotopologue distribution as indicated in each figure.

### Plasma itaconate pharmacokinetics

Itaconate kinetic data analysis was performed using the add-in program PKSolver 2.0 for Microsoft Excel^43^. Rat data depicted in Fig. 1 was generated through a non-compartmental analysis of plasma data after intravenous constant infusion input for 30 min with an infusion dose of 15 mg/kg/min itaconate. Mouse data depicted in Fig. 2 was calculated through non-compartmental analysis of plasma data after intravenous bolus input with an infusion dose of 400 mg/kg body weight itaconate.

### Statistics

Data visualization and statistical analyses were performed using GraphPad Prism (v9.2.0) and Adobe Illustrator CS6 (v.16.0.0). Figure 4j was created with BioRender.com. The type and number of replicates, number of animals (n), and the statistical test used are described in each figure legend. Data are presented as means ± s.e.m. or box (25^th^ to 75^th^ percentile with median line) and whiskers (min. to max. values). *P* values were calculated using a two-*t*-test, one-way *ANOV*A or two-way *ANOVA* with no correction for multiple comparisons and with *p* < 0.05; \*\**p* < 0.01; \*\*\**p* < 0.001, and ^#^*p* < 0.0001 as indicated in each figure legend. Data collection and analysis were performed blind to the conditions of the experiment.

## Acknowledgment

This study was supported, in part, by US National Institutes of Health (NIH) grant R01CA234245 (to C.M.M.), DoD CDMRP HT9425-23-1-0388 (to P.C.), and internal funds from the Helmholtz Centre for Infection Research (to T.C.) and Technische Universität Braunschweig (to T.C.). We thank the NIH Common Fund Metabolite Standards Synthesis Core (NHLBI Contract No. HHSN268201300022C) for providing isotopic labeled [U-^13^C_5_]itaconate.

## Availability of data

All of the data associated with the study are in the paper or Extended Data. Source data will be deposited in the repository platform of Technische Universität Braunschweig with a dedicated doi upon acceptance of this manuscript.

## Contributions

T.C. - conceptualization, project administration, formal analysis, supervision, funding acquisition, investigation, visualization, methodology, writing—original draft, writing—review and editing. C.M.M –funding acquisition, supervision, writing—review and editing. P.C. –funding acquisition, supervision.- investigation. A.L.- investigation. H.F.W. – visualization, investigation, writing — review and editing. B.D. – validation, writing—review and editing.

## Figure Legends

**Extended Data Figure 1: Plasma metabolite levels after itaconate infusion. a**, Timeline of itaconate infusion and sample collection with two infusion cycles. **b**, Kinetic parameters for the two infusion cycles. **c**, Plasma lactate levels. **d**, Relative plasma citrate level. **e-t**, Plasma metabolite levels over time of **e**, malate, **f**, isoleucine, **g**, leucine, **h**, valine, **i**, alanine, **j**, aspartate, **k**, glutamate, **l**, glycine, **m**, histidine, **n**, lysine, **o**, methionine, **p**, phenylalanine, **q**, proline, **r**, serine, **s**, threonine, and **t**, tyrosine. T_1/2_ -elimination halftime; C_max_ -maximal plasma concentration; AUC - area under the curve. Experiments were performed with n = 4 rats and itaconate was infused twice with 15 mg/kg/min for each 30 min indicated in grey. Data are presented as means ± s.e.m. obtained from n = 4 rats. Two-way *ANOVA* compared to 0 min with * *p* < 0.05; ** *p* < 0.01.

**Extended Data Figure 2: Itaconate influences abundances of TCA cycle intermediates in tissues. a**, Schematic depicting an experimental overview of ^13^C itaconate tracing in mice with time points of tissue and plasma collections. **b**, Kinetic parameters after itaconate injection. **c**, Heat map depicting foldchange of TCA cycle intermediate abundances 15 min after itaconate injection relative to control injection diverse tissues. **d**, Succinate levels in different tissues. **e-j**, Levels of pyruvate and TCA cycle intermediates in **e**, kidney, **f**, brain, **g**, lung, **h**, liver, **i**, spleen, and **j**, heart. T_1/2_ - elimination halftime; C_max_ - maximal plasma concentration; AUC - area under the curve. Mice were injected with 400 mg/kg body weight [U-^13^C_5_]itaconate. Data are presented as heatmap (c) or means ± s.e.m (d-j) with tissue samples at 0 min (n = 4), 15 min (n = 5), and 45 min (n = 4). One-way *ANOVA* (c) or two-way *ANOVA* (d-j) with * *p* < 0.05; ** *p* < 0.01, *** *p* < 0.001, ^#^ *p* < 0.0001.

**Extended Data Figure 3: Itaconate modulates branched - chain amino acid metabolism in tissues. a**, Plasma methylmalonate levels. **b**, Levels of valine, leucine, and isoleucine relative to 0 min itaconate. **c**, Levels of methylmalonate in different mouse tissues. Abundances of **d**, isoleucine, **e**, leucine, and **f**, valine relative to 0 min itaconate in different mouse tissues. Mice were injected with 400 mg/kg body weight [U-^13^C_5_]itaconate. Data are presented as means ± s.e.m (a) or box (25^th^ to 75^th^ percentile with median line) and whiskers (min. to max. values) (b-f). Plasma samples at 0 min (n = 4), 15 min (n = 9), 30 min (n = 4), and 45 min (n = 4); tissue samples at 0 min (n = 4), 15 min (n = 5), and 45 min (n = 4). One-way *ANOVA* compared to 0 min itaconate (a) or two-way *ANOVA* (b-f) with * *p* < 0.05; ** *p* < 0.01.

**Extended Data Figure 4: Fate of** ^**13**^**C itaconate in kidney tissue. a-k** Isotopologue distribution from [U-^13^C_5_]itaconate on **a**, itaconate, **b**, pyruvate, **c**, lactate, **d**, alanine, **e**, citrate, **f**, α- ketoglutarate, **g**, glutamate, **h**, succinate, **i**, fumarate, **j**, malate, and **k**, aspartate in kidney tissue. Schematic of [U-^13^C_5_]itaconate use for TCA cycle metabolism with open bars depicting ^12^C and closed circles ^13^C shown in the middle of the figure. Mice were injected with 400 mg/kg body weight [U-^13^C_5_]itaconate. Data are presented as means ± s.e.m with tissue samples at 0 min (n = 4), 15 min (n = 5), and 45 min (n = 4). Multiple unpaired *t*-test with * *p* < 0.05; ** *p* < 0.01, *** *p* < 0.001.

**Extended Data Figure 5: Itaconate labels citramalate and mesaconate in tissues and culture models. a**, Isotopologue distribution from [U-^13^C_5_]itaconate on itaconate, mesaconate, and citramalate in Huh7 cells cultured with 1 mM ^13^C itaconate for 24 h, **b**, Isotopologue distributions on metabolites from HepG2 cells cultured with 1 mM ^13^C itaconate for 24 h, and **c**, HEK-293 cells cultured with 3 mM ^13^C itaconate for 120 min. **d**, Label on metabolites in the urine. **e**, Chemical structures of itaconate, mesaconate, and citramalate. **f**, Labeling (1 – M0) on itaconate **g**, mesaconate, and **h**, citramalate from [U-^13^C_5_]itaconate in different tissues. Mice were injected with 400 mg/kg body weight [U-^13^C_5_]itaconate. Data are presented as means ± s.e.m. Plasma samples at 0 min (n = 4), 15 min (n = 9), 30 min (n = 4), and 45 min (n = 4); tissue samples at 0 min (n = 4), and 15 min (n = 5). Multiple unpaired *t*-test (f-h) with * *p* < 0.05; ** *p* < 0.01, *** *p* < 0.001, ^#^ *p* < 0.0001.

